# Stretching mucins: revealing the complex rheology of a natural glycoprotein network

**DOI:** 10.64898/2026.05.15.725541

**Authors:** Bianca Hazt, George D. Degen, Lucas Warwaruk, Daniel J. Read, Adam O’Connell, Oliver G. Harlen, Gareth H. McKinley, Anwesha Sarkar

## Abstract

Flow and extensional deformation of mucin networks are fundamental in mucus biophysics, governing how mucus functions as a protective and lubricating, and transport-facilitating layer. While the shear and oscillatory rheology of mucin solutions have been characterized in considerable detail, their behavior under extensional deformation remains comparatively understudied. Here, we report a concentration-dependent transition in extensional flow response of mucin solutions using a bespoke dripping-onto-substrate extensional rheometer. We show that mucin solutions at the lower concentrations undergo linear filament thinning, whereas semidilute mucin solutions form highly extensible filaments, with radius decaying exponentially in time, consistent with the elastocapillary thinning observed in solutions of high molecular weight synthetic polymers. Remarkably, at higher mucin concentrations inter-chain mucin associations produce a sudden reduction in the apparent elastocapillary relaxation time. We demonstrate how increasing macromolecular concentration redistributes the balance between viscous and elastic stresses during capillary thinning in a biopolymer network and reveal a concentration-driven reduction in mucin filament extensibility.

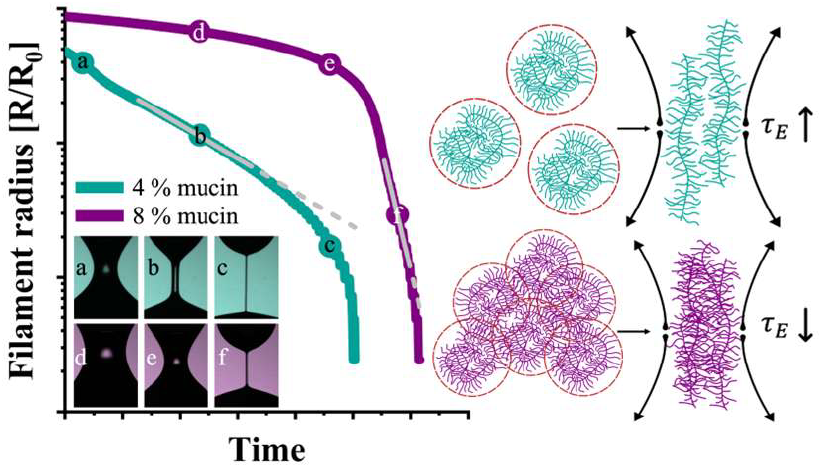

---

Mucus is a biological hydrogel that is essential for ocular, respiratory, gastrointestinal, and reproductive health. Its complex rheological behavior arises from both its composition and environment and directly impacts its ability to lubricate surfaces and facilitate clearance of undesirable particulates or microorganisms^1^. The material properties of mucus arise primarily from gel-forming mucin glycoproteins assembled into supramolecular disulfide-linked networks, and from their interactions with other minor mucosal components^2^. Variations in mucin concentration, which *in vivo* span nearly two orders of magnitude depending on the biological site and pathological state, may lead to distinct mechanical responses under physiological stresses. In different physiological settings, this tunability serves distinct (and sometimes antagonistic) roles. Highly elastic, weakly extensible mucus is required to form a robust cervical mucus plug during a healthy pregnancy^3, 4^, whereas in the airways excessively viscoelastic mucus can impair transport and clearance in diseases such as cystic fibrosis or chronic obstructive pulmonary disease^5, 6^. Concentration-dependent changes in the shear rheology of mucus and mucin solutions are well documented^7-10^, as well as in the dynamics of filament thinning for saliva^11^. However, how mucin concentration regulates the extensional response of mucin solutions, which is central to filament breakup, airway clearance, and transport through constricted geometries, has not been established.

Experimental methods such as capillary breakup extensional rheometry (CaBER)^12^ or the more recently developed dripping-onto-substrate (DoS) reometry^13^ provide a means to characterize the extensional properties of a wide range of low viscosity viscoelastic liquids that are only available in small test volumes. In DoS extensional rheometry, a hydrophilic partially-wetting substrate is brought into contact with a pendant drop. Upon contact, the drop rapidly wets the substrate, and an unstable liquid bridge is formed. Measurements of the time-evolving filament radius as the bridge thins under the action of capillarity are captured using high-speed imaging and used to determine an apparent transient extensional viscosity, as well as other rheological properties of the fluid, *e*.*g*., the dynamic viscosity and extensional relaxation time. Compared to CaBER, DoS rheometry avoids problems due to inertio-capillary oscillations in low-viscosity, weakly elastic materials, enabling measurements of weakly viscoelastic materials with sub-millisecond relaxation timescales^14^. Although DoS rheometry has been used to measure the extensional properties of synthetic polymer solutions such as polyethylene oxide^15^, and of biopolymeric systems including proteins^16^, polysaccharides^17, 18^, and native biological fluids such as saliva and bronchial sputum^19^, extensional measurements on reconstituted glycoprotein solutions remain scarce, with existing reports limited to unpurified gastric mucin-containing formulations examined at concentrations up to 4 wt%^20^.

In this letter, we investigate the rheology of a natural glycoprotein network, namely purified mucin solutions, over a range of concentrations spanning 0.25 to 8 wt%. With increasing concentration, a qualitative transition in the extensional rheological behavior is observed at concentrations corresponding to the formation of a viscoelastic gel. This transition is characterized by an increase in elasticity and dynamic viscosity that delays the formation of an extensional filament in DoS, as well as a reduction in the apparent relaxation time extracted from the elastocapillary regime. To our knowledge, this behavior has not previously been identified for mucus or mucin solutions.

The mucin used in this work was extracted from bovine submaxillary glands (BSM, Sigma Aldrich), a widely used and well-characterized model mucin system^7^. All mucin solutions were prepared in 10 mM 4-(2-Hydroxyethyl)piperazine-1-ethanesulfonic acid (HEPES) buffer at pH 7.0. To minimize contributions from non-mucin components, such as albumins and DNA, BSM solutions were purified by extensive dialysis against ultrapure water^21^. The resulting mucin exhibits a radius of gyration *R*_*g*_ = 165 nm and weight-average molecular weight 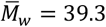 MDa, as previously determined by AF4-MALS^22^, yielding an estimated overlap concentration *c*^∗^ = 0.34 wt%^22^, and intrinsic viscosity [*η*] = 227 mL g^-1^ using the Graessley relation^23^.

We measured the linear viscoelastic moduli as a function of angular frequency using a strain-controlled ARES-G2 (TA Instruments, Newcastle, DE), equipped with a 2° 40 mm cone-and-plate geometry; interfacial effects and evaporation were avoided by using a 2 vol% Tween 80 layer and a mineral oil overlay, respectively, deposited at the air-liquid interface. The frequency dependence of the storage and loss moduli *G*′ (*ω*), *G*″ (*ω*) for 2, 4 and 5 wt% BSM shows a liquid-like linear viscoelastic response characteristic of the terminal regime, with *G*″ ∝ *ω* and *G*′ ∝ *ω*^2^, at frequencies below the inverse of the longest relaxation time. Although the measurements are restricted to this low-frequency limit, we can estimate a characteristic relaxation time by extrapolating the measured data up to the frequency where *G*′ = *G*″ (Figure 1). For BSM at 2 and 4 wt%, the inverse of the crossover frequencies yield relaxation times of ∼1.6 and ∼2.7 ms, respectively. A similar estimate for 5 wt% is provided in Figure S1 (Supporting Information), giving *τ* = 4.3 ms, consistent with unentangled viscoelastic behavior on the timescales probed. For truly dilute polymers (*c*/*c*^∗^ ≪ 1), the longest relaxation time approaches a constant value corresponding to the Zimm relaxation time at infinite dilution^24^, reflecting that polymer dynamics are governed by hydrodynamic interactions mediated by the solvent. This timescale can be estimated as 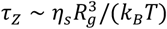, which, using the value of *R*_*g*_ determined from the Asymmetric Flow Field-Flow Fractionation with Multi-Angle Light Scattering (AF4-MALS) experiment, yields a value on the order of 1 ms. The systematic increase in the relaxation time with concentration we measure here is attributable to intermolecular interactions arising above the overlap concentration *c*^∗^. In this regime, chains are no longer isolated and relax within a progressively crowded medium, in which the effective resistance to motion increases with concentration due to the presence of surrounding polymers, leading to slower chain relaxation^25, 26^.

**Figure 1.**
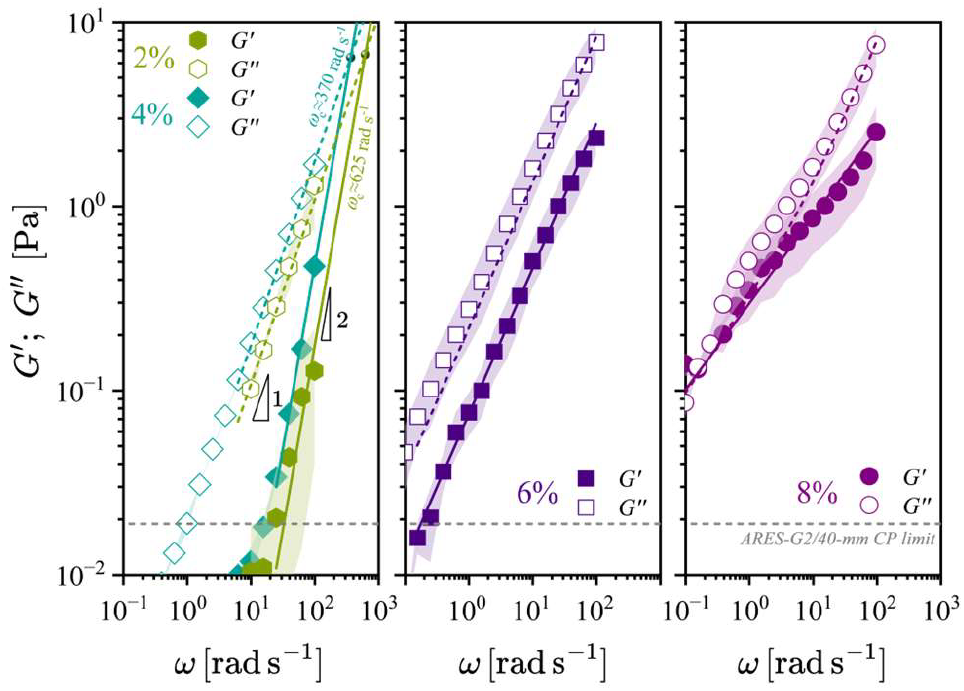
Dynamic viscoelastic response of BSM solutions obtained from small amplitude oscillatory shear frequency sweeps at 2, 4, 6, and 8 wt%. Mean values ± standard deviation (shaded regions) are shown for triplicate measurements. Solid and dashed lines represent fits to *G*′ and *G*”, respectively. Fits correspond to a Maxwell model (2 and 4 wt%), a Scott-Blair critical gel model with *β* ≈ 0.81 (6 wt%), and a fractional Kelvin-Voigt liquid (8 wt%; with fitted parameters 𝕍 = 0.054 Pa s^*α*^, 𝔾 = 0.396 Pa s^*β*^, *α* = 1, *β* = 0.47).

Figure 1 shows that further increasing the BSM concentration to 6 and 8 wt% leads to a pronounced change in the linear viscoelastic response. At 6 wt%, both moduli scale as common power laws with frequency, with a nearly constant ratio of *G*″/*G*′ over the measured frequency range, consistent with the Winter-Chambon criterion^27^. Fitting the Scott-Blair model for a critical gel to the 6 wt% data yields *β* ≈ 0.81, which determines a constant value of tan *δ* = *G*″/*G*′ = tan(*πβ*/2). At 8 wt% the elasticity increases further and the ratio tan *δ* now increases with frequency (see Figure S1, Supporting Information), as expected for a viscoelastic gel^27^. The emergence of this response at higher concentration suggests that elastic stresses arise from an increasing density of transient physical crosslinks between mucin macromolecules, which further constrain chain motion but without giving rise to an equilibrium elastic plateau modulus within the experimentally accessible timescales. A two-element, four-parameter fractional Kelvin-Voigt model has previously been employed to compactly describe the linear viscoelastic behavior of weakly viscoelastic biopolymer networks such as unpurified BSM^7^. Using this constitutive representation, we can identify an increasing elastic contribution as the mucin concentration is raised from 6 to 8 wt%; there is a decrease in the power law exponent (*β*) characterizing the concomitant frequency response of the viscoelastic network and an increase in the quasi-property (𝔾) characterizing the strength of the viscoelastic response (see Supporting Information, Tables S1 and S2, and Figure S1 for additional details). Consistent with the emergence of a strong concentration-dependent linear viscoelastic response, steady shear viscosity measurements reveal pronounced shear-thinning behavior for 8 wt% BSM (Figure S2a, Supporting Information). We can systematically demonstrate that the elasticity arises from intermolecular interactions (see Figure S2b, Supporting Information), as treatment with a typical chaotropic agent such as 8M urea (which disrupts hydrogen bonding) or 20 mM dithiothreitol (which reduces disulfide bonds) leads to progressive disruption of intermolecular interactions and reduction of the strength of the viscoelastic response.

The custom DoS experimental setup consisted of a syringe pump (PHD 2000, Harvard Apparatus) used to dispense discrete volumes at low flow rate through a blunt-end nozzle of radius *R*_0_ = 0.635 mm. A pendant drop was formed, and the substrate was brought into contact with the droplet at a fixed distance (*H*) below the nozzle, corresponding to an aspect ratio *AR* ≡ *H*/2*R*_0_ = 2.2. The necked region was imaged using a high-speed camera (NOVA S20, Photron) operating at an image acquisition rate of *f*_*s*_ = 16,500 Hz, equipped with a high-magnification zoom lens (LAOWA, Venus Optics), and a light emitting diode (SOLIS-525C, Thorlabs Inc.) to provide back-light illumination of the liquid filament. Here, the limiting filament capture rate, calculated as (*FCR*_0_ = *f*_*s*_ ln(*m*_0_),^14^ where *m*_0_ = *R*_0_/*R*_*res*_ = 430.46, and *R*_*res*_ = 1 pixel, is *FCR*_0_ ≈ 1.0 × 10^5^ s^-1^. The filament radius is a function of axial position and time, *R* = *R*(*z, t*). Following the image-analysis procedure described by Warwaruk *et al*.^14^, we take the measured radius as the axial minimum at each instant, *R*(*t*) = *min*_*z*_{*R*(*z, t*)}. Figures 2a-h show measurements of the BSM filament thinning at different concentrations as well as for the HEPES buffer alone. Replicated measurements for each sample are presented in Figure S3 (Supporting Information).

**Figure 2.**
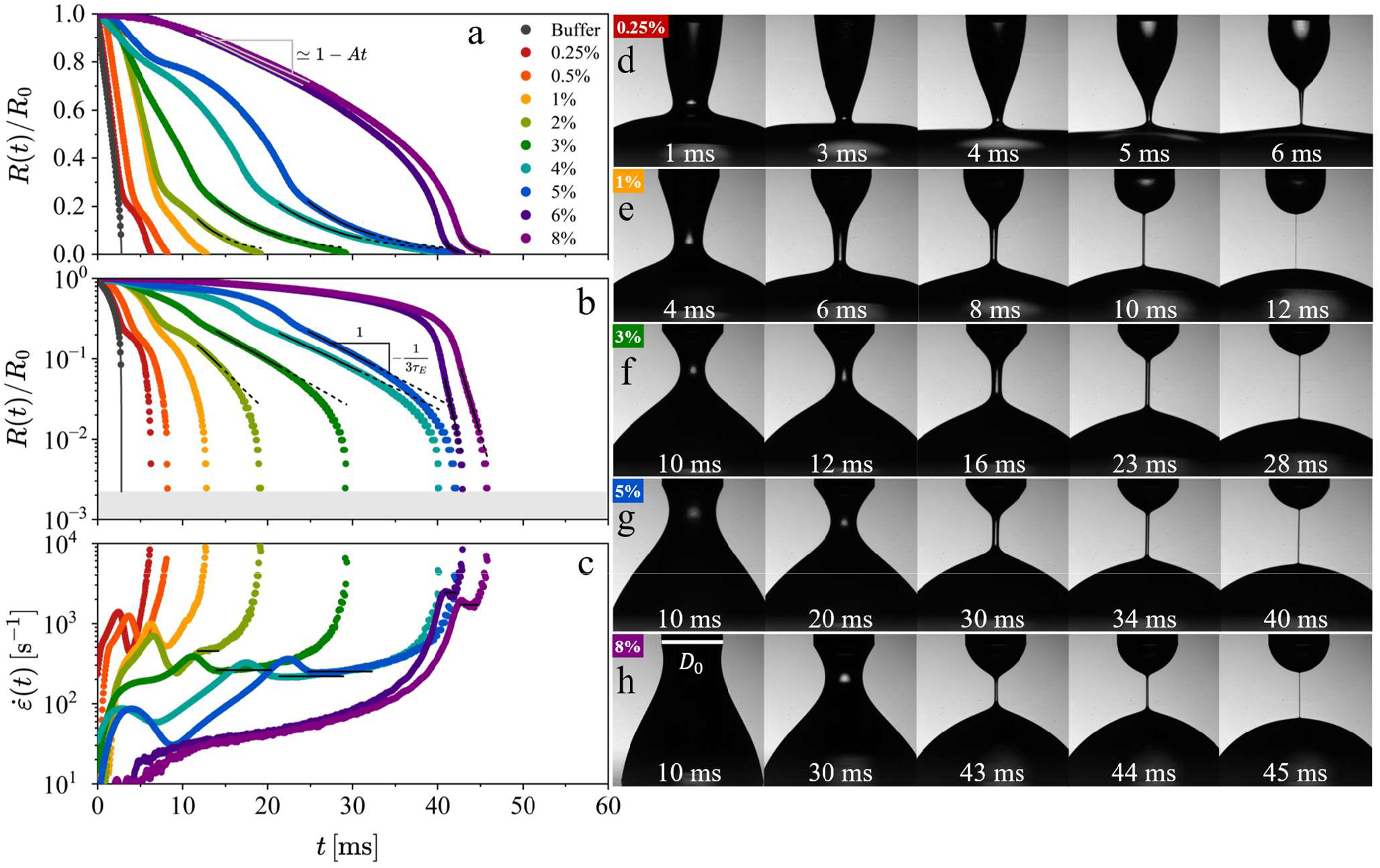
Temporal evolution of the normalized midpoint filament radius *R*(*t*)/*R*_0_ in capillary thinning for BSM solutions from 0.25 wt% to 8 wt%, and for 10 mM HEPES buffer at pH 7.0. (a) Data presented on a linear-linear axis highlight the overall thinning behavior across concentrations, with a VC balance established at intermediate times in the 6 and 8 wt% BSM solutions indicated by a linear fit. (b) The same profiles shown on semi-logarithmic axes (logarithmic ordinate, linear abscissa) reveal the emergence of an exponential decay regime at concentrations *c* ≥ 2 wt% characteristic of EC thinning. The gray shaded region indicates the minimum measurable radius *R*_*res*_ = 1.51 *µ*m, corresponding to one pixel. (c) The extensional rate 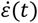 extracted from the radius evolution plotted as a function of time. For each plot, the time interval over which a constant extension rate is established [and from which the characteristic relaxation time is obtained] is marked with solid black lines and extended by black dashed lines. Representative high-speed image sequences illustrate the capillarity-driven filament thinning and final breakup of the BSM solutions for (d) 0.25 wt%, (e) 1 wt%, (f) 3 wt%, (g) 5 wt%, (h) 8 wt%. The scale bar in (h) represents the nozzle diameter *D*_0_ = 2*R*_0_ = 1.27 mm.

For a slender fluid filament undergoing capillarity-driven thinning, there are several distinct regimes depending on the balance between the capillary pressure, inertial, viscous, and elastic contributions to the stress. In the limit where viscous and elastic stresses are negligible, the thinning is governed by a balance between inertia and surface tension. In this inertio-capillary (IC) limit, the time-evolving minimum filament radius is given by *R*(*t*)/*R*_0_ = *α* [(*t*_*IC*_ − *t*)/*t*_*R*_]^2/3^, where, *t*_IC_ is the critical time to breakup, *α* is a system-dependent prefactor which can vary between 0.4 ≤ *α* ≤ 1,^16^ *R*_0_ is the nozzle radius, and *t*_*R*_ is the Rayleigh time, 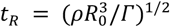, with surface tension *Γ* and the fluid density *ρ*. This expression accurately describes the thinning dynamics observed for the HEPES buffer. In Figure 2a,b, we show an IC fit with *α* ≈ 0.73, and in Figure S4 (Supporting Information) we show that the filament has an axially asymmetric and cone-shaped morphology prior to breakup^28^. Measured surface tension values can be found in the Supporting Information (Table S3, Figure S5). For more viscous Newtonian fluids, liquid filaments exhibit visco-capillary (VC) thinning with a minimum radius that decays linearly in time *R*(*t*)/ *R*_0_ = 1 − 0.0709(*t* − *t*_0_)/*t*_*VC*_, where *t*_*VC*_ = *ηR*_0_/*Γ* is the viscous timescale^29^. The separation between the IC and VC scalings is governed by the magnitude of the Ohnesorge number 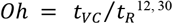. For small Ohnesorge numbers *Oh* ≪ 1, inertia dominates, and thinning follows IC scaling. Flows with large Ohnesorge numbers of *Oh* ≥ 1 are dominated by viscous stresses, and exhibit VC thinning^31, 32^. For our BSM solutions, the Ohnesorge number increases from *𝒪*(10^−3^) at the lowest concentration to *Oh* ≈ 0.78 at the highest concentration tested here (see Supporting Information, Table S4), indicating a transition from inertia-dominated to increasingly viscously dominated initial filament thinning conditions.

For viscoelastic fluids, the filament thinning may initially follow either IC or VC scalings (depending on the background viscosity) until elastic stresses become significant, after which the thinning dynamics shift towards an elasto-capillary (EC) balance^33^. In this regime, the minimum filament radius is predicted to decay exponentially with time:

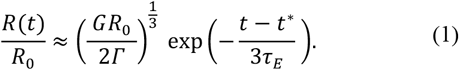

Here, *G* is the elastic modulus, *t*\* is the experimentally observed time at which the flow transitions to EC thinning and a cylindrical filament is formed, as is evident in the images shown in Fig 2(e)-(h). By fitting the observed exponential decay to Eq. (1), an effective extensional relaxation time *τ*_*E*_ can be extracted. This EC regime holds until the dissolved mucin chains reach their limit of finite extensibility and the filament then transitions to a terminal finitely extensible nonlinear elastic regime^15, 34, 35^.

For BSM concentrations of 0.25 (Figure 2d) and 0.5 wt%, the filament profile initially develops a cone-shaped neck characteristic of the IC regime before forming a short-lived, slender filament that undergoes capillary breakup before an axially uniform filament of constant radius can be established. While this transient response indicates the presence of elastic effects, the filament is too short-lived to establish a well-defined EC regime. By contrast, BSM concentrations between 1 and 5 wt% produce axially uniform filaments (Figure 2e, image at 10 ms) that undergo a period of exponential thinning consistent with eq. (1) (denoted by black lines in Figures 2a and 2b). This regime can also be seen in Figure 2c, which shows the time-evolution of the local extensional strain rate 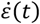 at the minimum radius

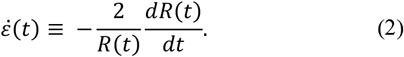

In the EC regime the extensional strain rate approaches a constant value of 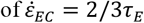. This is preceded by an inertioelastic overshoot as discussed by Tirtaatmadja *et al*.^36^ and Zinelis *et al*.^37^, characterized by a rapid increase in 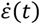 followed by a sharp decrease, associated with a coil-stretch transition in polyethylene oxide (PEO)^15^ and high-M_*w*_ polyacrylamides (PAM)^38^. For BSM concentrations between 2 and 5 wt% both the onset time and the duration of the exponential thinning regime increases with concentration consistent with the expected behavior of increasingly viscoelastic synthetic polymer solutions^39^.

The filament capture rate (FCR), defined as FCR = *f*_*s*_ ln(*m*),^14^ where *m* is the spatial dynamic range, is a figure of merit for filament thinning experiments that reflects the resolution of the EC regime and, equivalently, the largest extensional strain rate that can be reliably measured. Here, FCR varied between 67,700 and 74,400 s^-1^ for the imaging conditions used to visualize this concentration range (2-5 wt%), indicating that effective values of *τ*_*E*_ can be reliably extracted from the observations of EC regime. It is evident from Fig 2b and Fig 3b that for the 6 and 8 wt% BSM concentrations, the thinning of the filament is qualitatively different to that observed at lower concentrations. The extensional strain rate 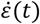 increases monotonically with time and the filament radius decreases approximately linearly in time for the first 30 ms, consistent with VC thinning. The subsequent elastocapillary thinning regime is markedly short-lived compared to lower concentrations. FCR values of 52,100 and 56,800 s^-1^ are obtained for 6 and 8 wt% BSM, respectively. Despite the smaller transition radius at the onset of the EC regime and the short period of time for which an elastocapillary balance is established, the filament capture rate for our DoS system is large enough to accurately resolve the thinning dynamics (Figure 2c, h) and determine the corresponding extensional relaxation time.

**Figure 3.**
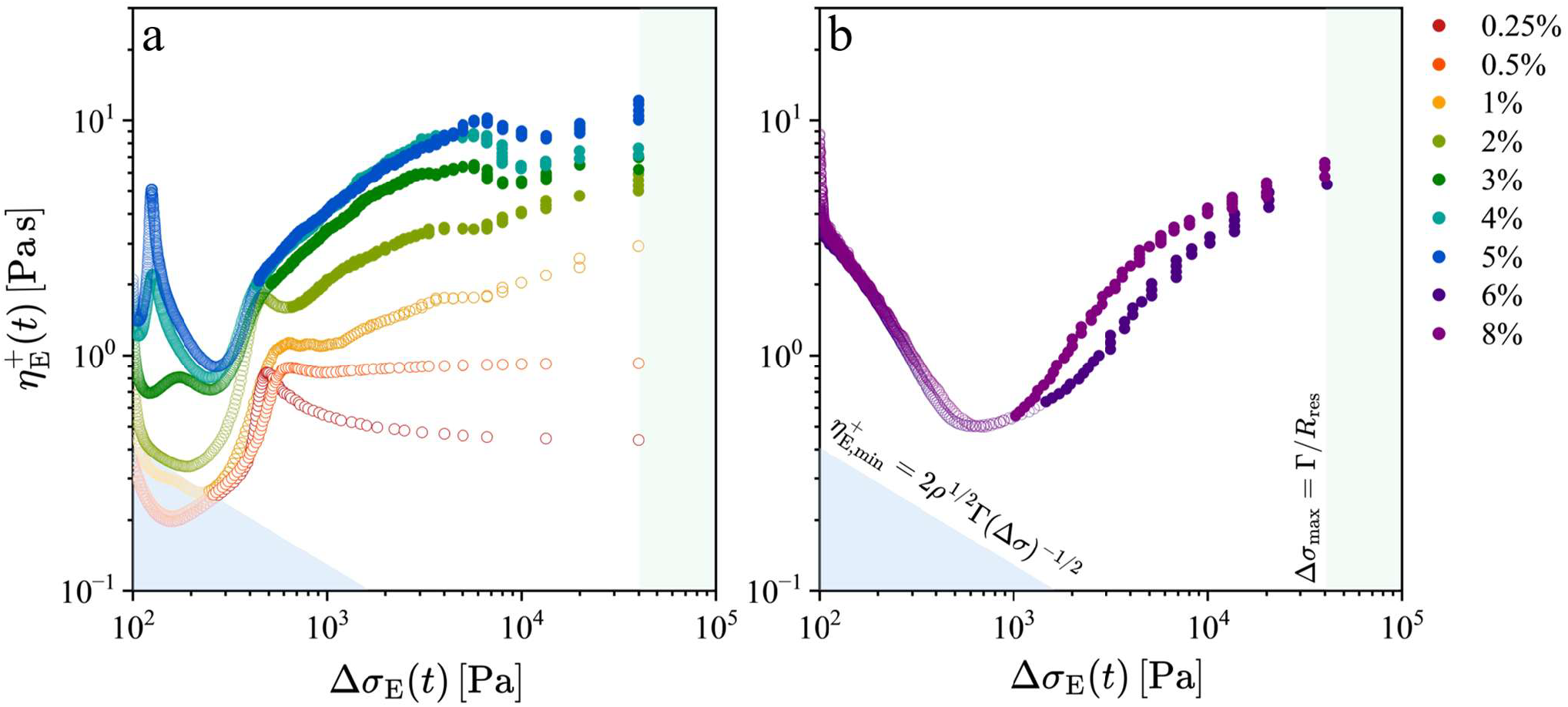
Apparent transient extensional viscosity as a function of the time-evolving extensional stress difference for BSM solutions: (a) 0.25-5 wt% and (b) 6 and 8 wt%. Open symbols denote pre-elastocapillary (EC) regimes; filled symbols denote data at and beyond EC thinning. The colored shaded regions denote areas in which measurements of 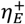 and Δ*σ*_*E*_ become unreliable; these corresponding limits are described in further detail in Figure S6 of the Supporting Information.

Agnostic of imposing model-specific filament thinning dynamics (e.g., VC or EC thinning), the apparent transient extensional viscosity 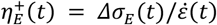 can always be estimated from sufficiently time-resolved measurements of the minimum filament radius *R*(*t*) for each mucin solution. Figure 3 summarizes the evolution of the transient extensional viscosity 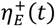 for the mucin solutions as a function of the time-evolving capillary pressure Δ*σ*_*E*_ (*t*) = *Γ*/*R*(*t*), which drives the thinning of the liquid filament. Shaded regions in Figure 3 correspond to limit lines in which measurements of 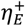 and Δ*σ*_*E*_ should not be interpreted, as the thinning dynamics become dominated by inertio-capillary effects rather than material contributions to the tensile stresses [corresponding to the constraint 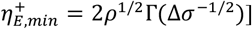. The resolution of the imaging system also sets a smallest measurable radius and an upper bound on the stress, Δ*σ*_*max*_ = Γ/*R*_*res*_. Detailed limit lines that constrain the measures of 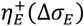 and facilitate interpretation of DoS measurements are further discussed by Warwaruk *et al*.^14^ and shown in Figure S6 (Supporting Information).

As the mucin filaments thin, the tensile stress difference increases and all samples exhibit strain stiffening within the EC regime (Figure 3, filled symbols), with 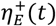 increasing approximately linearly with Δ*σ*_*E*_(*t*) on log-log axes for all concentrations, consistent with an elasto-capillary stress balance. For the more concentrated samples (6 and 8 wt%), an initial apparent extensional-thinning region appears before the EC transition at Δ*σ*_*E*_(*t*)∼10^3^Pa. The magnitude of 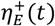 increases as expected from 1-5 wt% BSM but subsequently decreases at higher concentration (6-8 wt%) despite the larger tensile stresses achieved, indicating a change in the characteristic extensional relaxation dynamics.

The effective relaxation times, *τ*_*E*_, extracted from the measured thinning dynamics in the EC regime are shown in Figure 4 as a function of BSM concentration. For BSM concentrations up to 4 wt%, the relaxation times obtained from extrapolating to the cross-over frequency in Small-Amplitude Oscillatory Shear (SAOS) and from DoS extensional experiments agree within experimental uncertainty (see Supporting Information, Table S5). This correspondence suggests that, in this concentration range, the extensional responses of the BSM solutions are governed by the same longest relaxation time that characterizes their small-strain shear behavior. However, at 5 wt% the relaxation time obtained from DoS is significantly lower than the linear relaxation time (see Table S5, Supporting Information).

**Figure 4.**
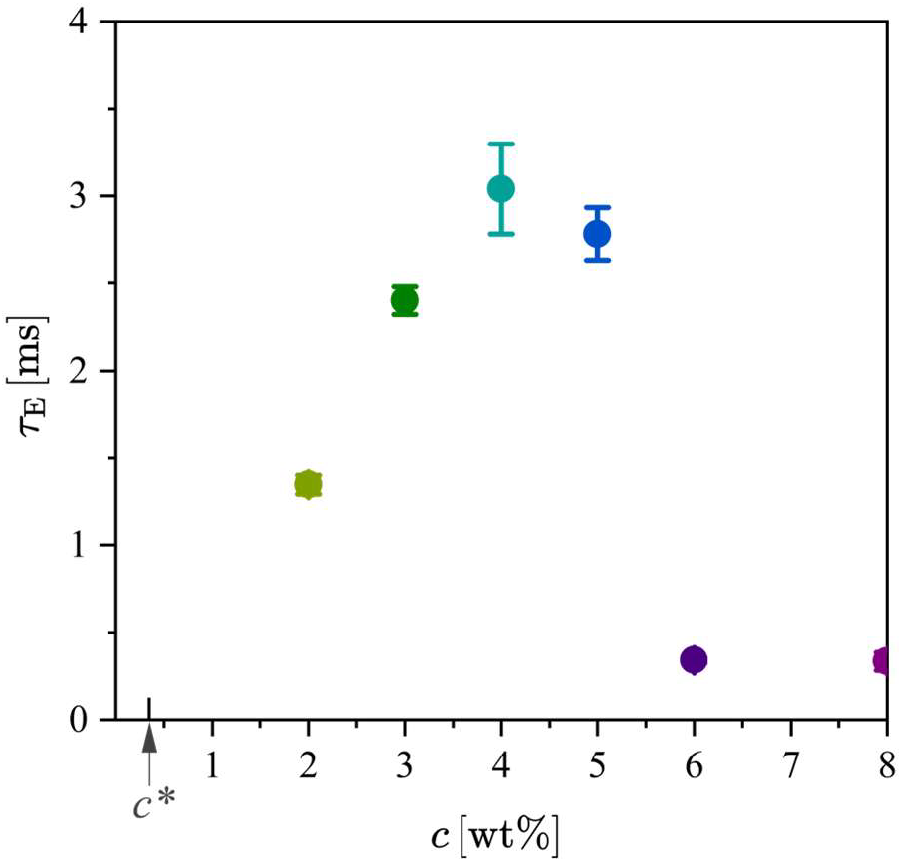
Extensional relaxation times extracted from the elastocapillary (EC) regime for each BSM concentration at which an EC regime was observed. The estimated overlap concentration *c*\* for BSM is indicated on the graph.

For higher BSM concentrations of 6 and 8 wt%, the relaxation times extracted from measurements in the EC regime decrease substantially below those measured for BSM concentrations *c* ≤ 5 wt%. This abrupt qualitative change in the filament thinning behavior has not, to our knowledge, been previously reported for mucin solutions. The potential physiological consequences of such behavior, particularly in the context of impaired mucus transport in health and disease, provide a direction for future studies. The relationship between the characteristic time extracted from EC thinning, *τ*_*E*_, and the linear viscoelastic relaxation time, *τ*, has been shown by Calabrese et al. (2024) to depend on macromolecular extensibility for model DNA^40^. For highly extensible molecules, *τ*_*E*_ ≈ *τ*, while for polymers with smaller extensibility a reduced ratio *τ*_*E*_ /*τ* is observed^40^. Here, we observe a precipitous decrease in *τ*_*E*_ as BSM concentration increases. While changes in concentration do not alter the intrinsic extensibility of the individual mucin chains, they do modify the bulk extensional flow response of the fluid filament and the condition under which elastic stresses become dominant. Numerical simulations of capillarity-driven thinning for entangled polymer systems described by the Doi-Edwards-Marrucci-Grizzuti (DEMG) and Rolie-Poly (RP) models predict a delayed transition to the EC regime compared to semi-dilute systems^41^. For these entangled systems, with increasing constraint density, the transition to the EC regime is expected to happen only close to the filament break time^41^. However, this happens gradually with increasing polymer concentration beyond *c*_*e*_ as entanglement effects become important and does not explain the precipitous transition observed here for this physically associated biopolymer network.

Consistent with observations in entangled polymer solutions^17, 41, 42^, the 6 and 8 wt% BSM samples exhibit a delayed onset of the EC regime, a reduced *τ*_*E*_, and no clear signature of a coil-stretch transition. However, unlike classical entangled polymer solutions, the linear viscoelastic response of these BSM samples does not display a well-defined plateau modulus but instead displays the signatures of a weakly elastic gel. This indicates that the underlying constraints are not purely topological entanglements. Rather, the concentrated BSM solutions are better described as transient, weakly elastic networks in which intermolecular associations impose constraints that limit filament extensibility. In this sense, while the DEMG/RP models attribute reduced extensibility to an increased number of entanglements per chain, a similar macroscopic response emerges here from transient physical associations, leading to a shortened elasto-capillary regime, lower molecular extensibility, and a reduction in the value of the extensional relaxation time *τ*_*E*_ extracted from analysis of the filament thinning dynamics.

To visualize concentration-dependent changes in the microstructure of the BSM samples, samples were chemically fixed with glutaraldehyde and subjected to critical point drying prior to imaging, following established protocols^43, 44^. At a BSM concentration of 2 wt%, no material remained adhered to the glass substrate after drying, indicating the absence of a fixed or percolated network. At intermediate concentrations (5 and 6 wt%, Figure S7, Supporting Information), some BSM remained attached to the glass substrate, suggesting the emergence of intermolecular associations, which allowed the visualization of the network. At higher concentrations (8 wt%), the dehydrated BSM remained attached to the substrate and exhibited a continuous, porous gel-like morphology (Figure 5), with multiscale connectivity evident throughout the structure. Although features such as the distribution of apparent pore sizes are likely influenced by the drying process and should therefore not be overinterpreted, the emergence of a visible, interconnected network at higher mucin concentration, which is absent in the less concentrated samples, is consistent with the onset of concentration-driven physical association and with the reduced extensional responses observed in the DoS measurements.

**Figure 5.**
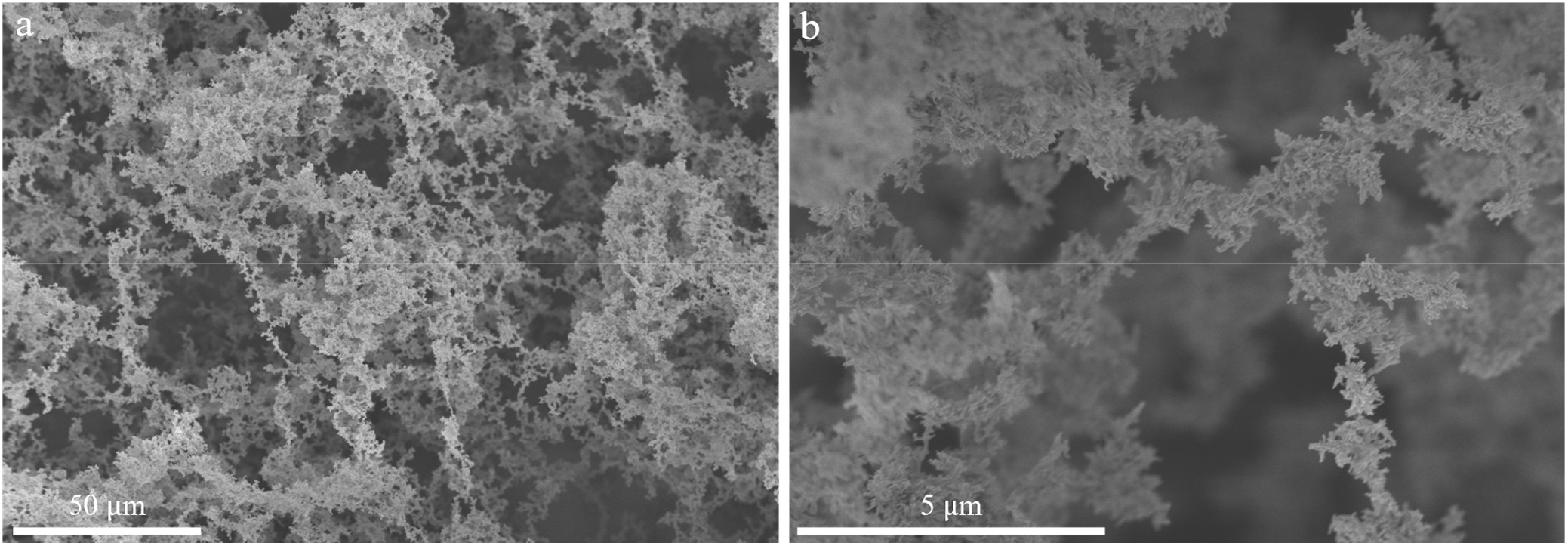
Representative Scanning Electron Microscopy (SEM) images obtained for an 8 wt% BSM sample. Scale bars represent (a) 50 *µ*m and (b) 5 *µ*m.

In conclusion, we have shown that both shear and the extensional response of mucin solutions are highly sensitive to concentration changes. Amid ongoing efforts to understand polymer interactions undergoing extensional flows^42^, these results position concentrated mucin solutions as a distinct class of weakly extensible, transiently interconnected networks. Semi-dilute mucin solutions form highly extensible filaments, whereas concentrated samples are dominated by viscous contributions to the tensile stress and enter the elastocapillary regime only at smaller filament radii than semidilute samples. The high filament capture rates (FCR > 52,100 s^-1^) of our DoS instrument provide very high spatial and temporal resolution, which enables the direct measurement of the extensional relaxation time *τ*_*E*_ for all semi-dilute and entangled mucin samples studied here. The solutions explored in this work ranging between 0.25 to 8 wt% in mucin concentration, span a crucial physiological and pathological spectrum, with direct relevance to airway mucus rheology for diagnosis of conditions where mucin overproduction (*e*.*g*., MUC5AC and MUC5B) drives disease severity. It should be noted that the purified mucin solutions in our study do not include the many minor mucosal components that are known to modify mucus rheology^2^. Further rheological studies systematically incorporating these components into model mucin systems will help bridge the gap towards understanding mucus behavior. We envision this work will contribute to new diagnostic tool strategies for assessing mucus rheology, and for guiding the design of bioinspired materials with tunable “stringiness”, *i*.*e*. resistance to capillarity-driven filament thinning mediated by concentration, solvent interactions and intermolecular interactions.

## ASSOCIATED CONTENT

### Supporting Information

The Supporting Information is available free of charge on the ACS Publications website.

Mathematical details of fractional rheological models; SAOS characteri**z**ation and viscoelastic model fits across concentrations, including tan *δ* analysis (Fig. S1); steady shear and chemical perturbation effects (Fig. S2); filament thinning dynamics and Bond number effects (Fig. S3**-**S4); surface tension measurement; extensional viscosity–stress relationships with limit lines (Fig. S5); SEM images for BSM at 5 and 6 wt% (Fig. S6); Ohnesorge number, surface tension and relaxation time comparisons (Tables S3**-**S5) (pdf)

## AUTHOR INFORMATION

### Author Contributions

The manuscript was written through contributions of all authors. All authors have given approval to the final version of the manuscript.

## ACKNOWLEDGMENTS

The authors gratefully acknowledge the Engineering and Physical Sciences Research Council (EPSRC) funded Centre for Doctoral Training in Soft Matter for Formulation and Industrial Innovation (SOFI^2^), Grant Ref. No. EP/S023631/1 for the financial support provided. This work was co-funded by Reckitt Benckiser Group, Plc. Author BH acknowledges the mobility funding from EPSRC to perform experiments during her research visit at the Massachusetts Institute of Technology (MIT), Cambridge, USA. The views expressed in this manuscript are those of the authors and do not necessarily reflect the position or policy of Reckitt Benckiser Group, Plc. The authors gratefully acknowledge Dr. Jim Bales (Edgerton Center, Massachusetts Institute of Technology) for providing access to the high-speed camera. Finally, we express our gratitude to Martin Fuller from the Astbury Biostructure Laboratory and Macauley Hough from the Leeds Electron Microscopy and Spectroscopy (LEMAS) for their help in Scanning Electron Microscopy (SEM) sample preparation and imaging.

## REFERENCES

(1) McShane, A.; Bath, J.; Jaramillo, A. M.; Ridley, C.; Walsh, A. A.; Evans, C. M.; Thornton, D. J.; Ribbeck, K. Mucus. Curr Biol 2021, 31 (15), R938–R945. DOI: 10.1016/j.cub.2021.06.093.

(2) Meldrum, O. W.; Yakubov, G. E.; Bonilla, M. R.; Deshmukh, O.; McGuckin, M. A.; Gidley, M. J. Mucin gel assembly is controlled by a collective action of non-mucin proteins, disulfide bridges, Ca(2+)-mediated links, and hydrogen bonding. Sci Rep 2018, 8 (1), 5802. DOI: 10.1038/s41598-018-24223-3.

(3) Critchfield, A. S.; Yao, G.; Jaishankar, A.; Friedlander, R. S.; Lieleg, O.; Doyle, P. S.; McKinley, G. H.; House, M.; Ribbeck, K. Cervical mucus properties stratify risk for preterm birth. PLoS One 2013, 8 (8), e69528. DOI: 10.1371/journal.pone.0069528.

(4) Smith-Dupont, K. B.; Wagner, C. E.; Witten, J.; Conroy, K.; Rudoltz, H.; Pagidas, K.; Snegovskikh, V.; House, M.; Ribbeck, K. Probing the potential of mucus permeability to signify preterm birth risk. Sci Rep 2017, 7 (1), 10302. DOI: 10.1038/s41598-017-08057-z.

(5) Hill, D. B.; Vasquez, P. A.; Mellnik, J.; McKinley, S. A.; Vose, A.; Mu, F.; Henderson, A. G.; Donaldson, S. H.; Alexis, N. E.; Boucher, R. C.; et al. A biophysical basis for mucus solids concentration as a candidate biomarker for airways disease. PLoS One 2014, 9 (2), e87681. DOI: 10.1371/journal.pone.0087681.

(6) Henderson, A. G.; Ehre, C.; Button, B.; Abdullah, L. H.; Cai, L. H.; Leigh, M. W.; DeMaria, G. C.; Matsui, H.; Donaldson, S. H.; Davis, C. W.; et al. Cystic fibrosis airway secretions exhibit mucin hyperconcentration and increased osmotic pressure. J Clin Invest 2014, 124 (7), 3047–3060. DOI: 10.1172/JCI73469.

(7) Rulff, H.; Schmidt, R. F.; Wei, L. F.; Fentker, K.; Kerkhoff, Y.; Mertins, P.; Mall, M. A.; Lauster, D.; Gradzielski, M. Comprehensive Characterization of the Viscoelastic Properties of Bovine Submaxillary Mucin (BSM) Hydrogels and the Effect of Additives. Biomacromolecules 2024. DOI: 10.1021/acs.biomac.4c00153.

(8) Kramer, C.; Rulff, H.; Ziegler, J. F.; Monch, P. W.; Alzain, N.; Addante, A.; Kuppe, A.; Timm, S.; Schrade, P.; Bischoff, P.; et al. Ileal mucus viscoelastic properties differ in Crohn’s disease. Mucosal Immunol 2024, 17 (4), 713–722. DOI: 10.1016/j.mucimm.2024.05.002.

(9) Wagner, C. E.; Krupkin, M.; Smith-Dupont, K. B.; Wu, C. M.; Bustos, N. A.; Witten, J.; Ribbeck, K. Comparison of Physicochemical Properties of Native Mucus and Reconstituted Mucin Gels. Biomacromolecules 2023, 24 (2), 628–639. DOI: 10.1021/acs.biomac.2c01016.

(10) Hazt, B.; Read, D. J.; Harlen, O. G.; Poon, W. C. K.; O’Connell, A.; Sarkar, A. Mucoadhesion across scales: Towards the design of protein-based adhesives. Adv Colloid Interface Sci 2024, 334, 103322. DOI: 10.1016/j.cis.2024.103322.

(11) Wagner, C. E.; McKinley, G. H. Age-dependent capillary thinning dynamics of physically-associated salivary mucin networks. Journal of Rheology 2017, 61 (6), 1309–1326. DOI: 10.1122/1.4997598.

(12) Rodd, L. E.; Scott, T. P.; Cooper-White, J. J.; McKinley, G. H. Capillary Break-up Rheometry of Low-Viscosity Elastic Fluids. Applied Rheology 2005, 15 (1), 12–27. DOI: 10.1515/arh-2005-0001.

(13) Dinic, J.; Zhang, Y.; Jimenez, L. N.; Sharma, V. Extensional Relaxation Times of Dilute, Aqueous Polymer Solutions. ACS Macro Lett 2015, 4 (7), 804–808. DOI: 10.1021/acsmacrolett.5b00393.

(14) Warwaruk, L.; Zinelis, K.; Ewoldt, R. H.; Macosko, C. W.; McKinley, G. H. DoS Dos and Don’ts. Arxiv 2025. DOI: 10.48550/arXiv.2511.17360.

(15) Dinic, J.; Sharma, V. Flexibility, Extensibility, and Ratio of Kuhn Length to Packing Length Govern the Pinching Dynamics, Coil-Stretch Transition, and Rheology of Polymer Solutions. Macromolecules 2020, 53 (12), 4821–4835. DOI: 10.1021/acs.macromol.0c00076.

(16) Lauser, K. T.; Rueter, A. L.; Calabrese, M. A. Small-volume extensional rheology of concentrated protein and protein-excipient solutions. Soft Matter 2021, 17 (42), 9624–9635. DOI: 10.1039/d1sm01253c.

(17) Dinic, J.; Sharma, V. Power Laws Dominate Shear and Extensional Rheology Response and Capillarity-Driven Pinching Dynamics of Entangled Hydroxyethyl Cellulose (HEC) Solutions. Macromolecules 2020, 53 (9), 3424–3437. DOI: 10.1021/acs.macromol.0c00077.

(18) Jimenez, L. N.; Martínez Narváez, C. D. V.; Sharma, V. Solvent Properties Influence the Rheology and Pinching Dynamics of Polyelectrolyte Solutions: Thickening the Pot with Glycerol and Cellulose Gum. Macromolecules 2022, 55 (18), 8117–8132. DOI: 10.1021/acs.macromol.2c00170.

(19) Bazilevsky, A. V.; Entov, V. M.; Rozhkov, A. N. Breakup of a liquid bridge as a method of rheological testing of biological fluids. Fluid Dynamics 2011, 46 (4), 613–622. DOI: 10.1134/s0015462811040119.

(20) Ahmad, M.; Ritzoulis, C.; Chen, J. Shear and extensional rheological characterisation of mucin solutions. Colloids Surf B Biointerfaces 2018, 171, 614–621. DOI: 10.1016/j.colsurfb.2018.07.075.

(21) Xu, F.; Liamas, E.; Bryant, M.; Adedeji, A. F.; Andablo-Reyes, E.; Castronovo, M.; Ettelaie, R.; Charpentier, T. V. J.; Sarkar, A. A Self-Assembled Binary Protein Model Explains High-Performance Salivary Lubrication from Macro to Nanoscale. Advanced Materials Interfaces 2019, 7 (1). DOI: 10.1002/admi.201901549.

(22) Hazt, B.; Read, D. J.; Harlen, O. G.; Poon, W. C. K.; O’Connell, A.; Connell, S. D.; Sarkar, A. Heat-Induced Structural Changes in Lactoferrin for Enhanced Mucoadhesion. ACS Appl Bio Mater 2025, 8 (11), 10255–10271. DOI: 10.1021/acsabm.5c01534.

(23) Graessley, W. W. Polymer chain dimensions and the dependence of viscoelastic properties on concentration, molecular weight and solvent power. Polymer 1980, 21 (3), 258–262. DOI: 10.1016/0032-3861(80)90266-9.

(24) Zimm, B. H. Dynamics of Polymer Molecules in Dilute Solution: Viscoelasticity, Flow Birefringence and Dielectric Loss. The Journal of Chemical Physics 1956, 24 (2), 269–278. DOI: 10.1063/1.1742462.

(25) Colby, R. H. Structure and linear viscoelasticity of flexible polymer solutions: comparison of polyelectrolyte and neutral polymer solutions. Rheologica Acta 2009, 49 (5), 425–442. DOI: 10.1007/s00397-009-0413-5.

(26) Rubinstein, M.; Colby, R. H. Polymer Physics; Oxford University Press, 2003. DOI: 10.1093/oso/9780198520597.001.0001.

(27) Winter, H. H.; Chambon, F. Analysis of Linear Viscoelasticity of a Crosslinking Polymer at the Gel Point. Journal of Rheology 1986, 30 (2), 367–382. DOI: 10.1122/1.549853 (accessed 4/22/2026).

(28) Day, R. F.; Hinch, E. J.; Lister, J. R. Self-Similar Capillary Pinchoff of an Inviscid Fluid. Physical Review Letters 1998, 80 (4). DOI: 10.1103/PhysRevLett.80.704.

(29) McKinley, G. H.; Tripathi, A. How to extract the Newtonian viscosity from capillary breakup measurements in a filament rheometer. Journal of Rheology 2000, 44 (3), 653–670. DOI: 10.1122/1.551105.

(30) Fardin, M. A.; Hautefeuille, M.; Sharma, V. Spreading, pinching, and coalescence: the Ohnesorge units. Soft Matter 2022, 18 (17), 3291–3303. DOI: 10.1039/d2sm00069e.

(31) Eggers, J. Nonlinear dynamics and breakup of free-surface flows. Reviews of Modern Physics 1997, 69 (3), 865–930. DOI: 10.1103/RevModPhys.69.865.

(32) Campo-Deaño, L.; Clasen, C. The slow retraction method (SRM) for the determination of ultra-short relaxation times in capillary breakup extensional rheometry experiments. Journal of Non-Newtonian Fluid Mechanics 2010, 165 (23), 1688–1699. DOI: 10.1016/j.jnnfm.2010.09.007.

(33) Dinic, J.; Sharma, V. Macromolecular relaxation, strain, and extensibility determine elastocapillary thinning and extensional viscosity of polymer solutions. PNAS 2019, 116 (8), 8766–8774. DOI: 10.1073/pnas.1820277116.

(34) Entov, V. M.; Hinch, E. J. Effect of a spectrum of relaxation times on the capillary thinning of a filament of elastic liquid. Journal of Non-Newtonian Fluid Mechanics 1997, 72 (1), 31–53. DOI: 10.1016/S0377-0257(97)00022-0.

(35) McKinley, G. H. Visco-elasto-capillary thinning and break-up of complex fluids. Rheology Reviews 2005.

(36) Tirtaatmadja, V.; McKinley, G. H.; Cooper-White, J. J. Drop formation and breakup of low viscosity elastic fluids: Effects of molecular weight and concentration. Physics of Fluids 2006, 18 (4). DOI: 10.1063/1.2190469.

(37) Zinelis, K.; Abadie, T.; McKinley, G. H.; Matar, O. K. The fluid dynamics of a viscoelastic fluid dripping onto a substrate. Soft Matter 2024, 20 (41), 8198–8214. DOI: 10.1039/d4sm00406j.

(38) Soetrisno, D. D.; Martínez Narváez, C. D. V.; Sharma, V.; Conrad, J. C. Concentration Regimes for Extensional Relaxation Times of Unentangled Polymer Solutions. Macromolecules 2023, 56 (13), 4919–4928. DOI: 10.1021/acs.macromol.3c00097.

(39) Rajesh, S.; Thievenaz, V.; Sauret, A. Transition to the viscoelastic regime in the thinning of polymer solutions. Soft Matter 2022, 18 (16), 3147–3156. DOI: 10.1039/d2sm00202g.

(40) Calabrese, V.; Shen, A. Q.; Haward, S. J. How Do Polymers Stretch in Capillary-Driven Extensional Flows? Macromolecules 2024, 57 (20), 9668–9676. DOI: 10.1021/acs.macromol.4c01604.

(41) Du, J.; Ohtani, H.; Kiziltas, A.; Ellwood, K.; McKinley, G. H. Capillarity-Driven Thinning Dynamics of Entangled Polymer Solutions. Macromolecules 2025, 58 (14), 7161–7177. DOI: 10.1021/acs.macromol.5c00782.

(42) Calabrese, V. Emergent Interpolymer Interactions in Flowing Polymer Solutions. ACS Macro Lett 2026. DOI: 10.1021/acsmacrolett.6c00010.

(43) Carpenter, J.; Wang, Y.; Gupta, R.; Li, Y.; Haridass, P.; Subramani, D. B.; Reidel, B.; Morton, L.; Ridley, C.; O’Neal, W. K.; et al. Assembly and organization of the N-terminal region of mucin MUC5AC: Indications for structural and functional distinction from MUC5B. PNAS 2021, 118 (39). DOI: 10.1073/pnas.2104490118.

(44) Nordgard, C. T.; Nonstad, U.; Olderoy, M. O.; Espevik, T.; Draget, K. I. Alterations in mucus barrier function and matrix structure induced by guluronate oligomers. Biomacromolecules 2014, 15 (6), 2294–2300. DOI: 10.1021/bm500464b.

