## Supplementary file for "Stretching mucins: revealing the complex rheology of a natural glycoprotein network"

List of Figures

Fractional rheological models……………………………………………………………………..…..4

Table S1 — Summary of the Maxwell and fractional mechanical models used in this work, their constitutive relations, characteristic relaxation timescale, and values of $\tan\delta$ from the oscillatory frequency response……………………………………………………………………………………….....4

Table S2 — Fitted quasi-properties ($\mathbb{V}$, $\mathbb{G}$) and power law exponents ($\alpha$, $\beta$) of the fractional Kelvin-Voigt Liquid model for BSM samples at 6 and 8 wt%…………………………………………………………..4

Figure S1 — (a) Linear viscoelastic response of BSM at 2, 4, and 5 wt%. Extrapolation of $G'' \sim \omega$ and $G' \sim \omega^2$ for BSM at 5 wt% yields a $G' = G''$ crossover at 232 rad s$^{-1}$ ($\tau \simeq 0.0043$ s). (b) Fractional Kelvin-Voigt liquid (FKV-Liquid) model fit for 6 and 8 wt% BSM (fitted parameters are reported in Table S2). (c,d) Frequency dependence of $tan\,\delta$. Solid lines represent a single-mode Maxwell fit where $tan\,\delta = 1/(\tau\omega)$ for 4 and 5 wt% BSM, a critical gel fit for BSM at 6 wt%, where $tan\,\delta = tan(\pi\beta/2) = G''/G'$, and a FKV liquid model fit for 8 wt%, where $tan\,\delta = [\mathbb{V}\omega + \mathbb{G}\omega^{\beta}\,sin(\pi\beta/2)]/[\mathbb{G}\omega^{\beta}\,cos(\pi\beta/2)]$. Panel (c) is shown in semilogarithmic representation (logarithmic x-axis), while panel (d) is plotted on log-log axes……………………..……….6

Figure S2 — (a) Steady-shear viscosity as a function of shear rate, measured using a 40 mm cone-and-plate geometry (2° CP-40 mm). The minimum rheometer sensitivity is shown as a grey dashed line. BSM samples at 8, 4, 2, 1, and 0.5 wt% were prepared in 10 mM HEPES buffer at pH 7.0. (b) Viscoelastic behavior of 8 wt% BSM in the absence or in the presence of 8 M urea, or 20 mM dithiothreitol. Shaded regions represent the standard deviations of three independent measurements. In the presence of DTT, which reduces disulfide bonds between mucin subunits, the covalent network at 8 wt% is disrupted, restoring a terminal liquid-like response with $G' {\sim} \omega^2$ and $G'' {\sim} \omega$, consistent with the Maxwell description. The corresponding relaxation time is $\tau = 8.60$ ms. In contrast, 8 wt% BSM in the presence of 8 M urea, which perturbs hydrogen bonding and weak non-covalent associations without reducing disulfide linkages, does not recover terminal behaviour of a viscoelastic liquid: while $G'' {\sim} \omega$ is preserved, the elastic modulus $G'$ exhibits an apparent $G' {\sim} \omega^{1.5}$ scaling over the accessible frequency window. (c) Steady-shear viscosity of the untreated 8 wt% BSM sample shown as a function of shear rate, compared with the complex viscosity extracted from $G'$ and $G''$, shown as a function of angular frequency, for 8 wt% BSM in the presence of either 8 M urea or 20 mM DTT……….…………………7

Figure S3 — Plots of the normalized filament radius $R(t)/R_0$ as a function of $(t_b - t)$, where $t_b$ is the time to breakup. For each concentration, at least three independent videos are included. In the terminal stage of the filament lifetime, replicates at a given concentration superimpose, and the trajectories for all concentrations converge as the filament approaches breakup. For completeness, data are presented using (a) log-linear, (b) log-log, and (c) linear-linear axes. The inset in (c) shows the terminal region on linear-linear axes………………...…………..……...…………………………………………...……………………………8

Terminal capillarity-driven thinning behavior of dilute BSM samples…………….………..……………..9

Figure S4 — Dripping-on-substrate measurements with $D_0 = 2R_0 = 1.27$ mm (a) for the solvent used in this work [10 mM HEPES buffer at pH 7.0 (blue)]. The data for 10 mM HEPES buffer at pH 7.0 are shown for the profile extracted from images performed at a Bond number $Bo = 0.35$. Snapshots of 10 mM HEPES buffer at pH 7.0 at a separation distance that yield Bond numbers of (b) $Bo = 0.51$ and (c) $Bo = 0.35$. The conical shape is most clearly observed at smaller needle-substrate separation distances $H$, corresponding to lower Bond numbers ($Bo = R_0 H\rho g/\Gamma$), as gravitational effects are reduced relative to capillarity at the moment the pendant drop touches the substrate. For the buffer, the Ohnesorge number $Oh = \eta_0/\sqrt{\rho\Gamma R_0}$ is $Oh \approx 4.4 \times 10^{-3}$, placing the system well within the inertio-capillary regime. The time to breakup ($t_b$) for HEPES buffer is 2.78 ms. (d) Normalized minimum filament radius $R(t)/R_0$ as a function of $t_b - t$ for 0.25 and 0.5 wt% BSM. In the final stage of thinning, both samples approach $R(t)/R_0 {\sim} (t_b - t\,)^1$ as indicated by the slope $n = 1$, consistent with a viscosity-dominated terminal thinning regime. (e-f) Corresponding high-speed image sequences for (e) 0.25 wt% ($t_b = 6.24$ ms) and (f) 0.5% BSM ($t_b = 8.11$ ms)……………..…..10

Figure S5 — Surface tension (Γ, mN m$^{-1}$) of BSM at 2 wt% as a function of time (s). Mean ± standard deviation from triplicate measurements is shown. Across all concentrations (0.25-8 wt%), the initial surface tension is indistinguishable within experimental resolution. Deviations only emerge at longer timescales, beyond the temporal window relevant to DoS measurements (< 1 s). Inset: variation of surface tension over 1-4 s, highlighting negligible early-time variations.....11

Figure S6 — Apparent transient extensional viscosity plotted as a function of the time-evolving extensional stress difference for BSM dispersions spanning 1 to 8 wt%. Limit lines, as discussed in reference 13 of the manuscript, are drawn using $\rho = 1080$ kg m$^{-3}$, $\Gamma = 61.7$ mN m$^{-1}$, $R_0 = 0.635$ mm, $R_{res} = 1$ pixel $= 1.51 \times 10^{-6}$ m, $FCR_0 = f_s \ln m_0$, with $f_s = 16500$ s$^{-1}$ and $m_0 = R_0/R_{res} = 430.46$. The horizontal lines shown in (a) indicate the asymptotic transient extensional viscosities $\eta_{E,\infty}$, as previously discussed (see the discussion on page S-8).....12

Figure S7 — Representative Scanning Electron Microscopy images of BSM samples prepared at (a) 6 wt% and (b) 5 wt%. Scales bars represent 10 µm.....13

Table S3 — Measured surface tension using a pendant drop tensiometer (OCA 25, DataPhysics Instruments, Germany). Drops (22 µL) were formed at the tip of a 1.65 mm DS500 GT syringe, and the liquid-air surface tension was obtained by fitting the drop profile to the Young-Laplace equation using dpiMAX software. A density of 1080 kg m$^{-3}$ was assumed in the fitting procedure. Measurements were performed in triplicate for each concentration; the reported value include the standard deviation. The initial surface tension was used in all capillarity-driven thinning analyses, as filament thinning occurs on timescales much shorter than those required for interfacial equilibration.....14

### Fractional rheological models

As discussed in the main text, the linear viscoelastic response of BSM samples at concentrations up to 5 wt% can be fitted to a Maxwell model, which consists of a linear mechanical combination of a Hookean spring and a dashpot in series. In this model, the resulting viscoelastic stress relaxation is governed by a single timescale $\tau$, which can be obtained from the crossover between power-law slopes of 2 and 1 for the storage and loss moduli, respectively[1]. Table S1 summarises the constitutive relation for the Maxwell model, the expected relaxation time and $\tan\delta$. Table S1 also shows the corresponding equations for a spring-pot (also known as the Scott-Blair fractional mechanical model) that describes a power-law response at the critical gel point, with a quasi-property $\mathbb{G}$ and power-law exponent $\beta$. Building from the spring-pot definition, the Fractional Kelvin-Voigt Model (FKVM) consists of two spring-pot elements arranged in a parallel configuration, with quasi-properties $\mathbb{V}$ and $\mathbb{G}$, and exponents $\alpha$ and $\beta$. The stress evolution for the FKV model is therefore represented by the expression

$$\sigma(t) = \mathbb{V}\frac{\mathrm{d}^{\alpha}\gamma(t)}{\mathrm{d}t^{\alpha}} + \mathbb{G}\frac{\mathrm{d}^{\beta}\gamma(t)}{\mathrm{d}t^{\beta}}.$$

Here, the 8 wt% BSM behavior is described by a limiting case of the FKVM in which $\alpha = 1$. In this scenario, the quasi-property $\mathbb{V}$ reduces to a viscosity $\eta$, and the configuration corresponds to a Fractional Kelvin-Voigt Liquid/Dashpot Model (FKV-Liquid) (Table S1). The fitted parameters for the 6 and 8 wt% BSM samples using the FKV-Liquid Model are summarised in Table S2.

Table S1 — Summary of the Maxwell and fractional mechanical models used in this work, their constitutive relations, characteristic relaxation timescale, and values of $\tan\delta$ from the oscillatory frequency response.

| | Constitutive relation | Relaxation timescale $\tau$ | $\tan\delta$ |
|---|---|---|---|
| Maxwell model 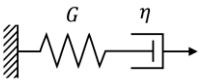  | $\sigma(t) + \frac{\eta}{G}\frac{\mathrm{d}\sigma(t)}{\mathrm{d}t} = \eta\frac{\mathrm{d}\gamma(t)}{\mathrm{d}t}$ | $\tau = \frac{\eta}{G}$ | $\tan\delta = \frac{G''}{G'} = \frac{1}{\tau\omega}$ |
| Spring-pot 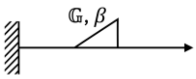  | $\sigma(t) = \mathbb{G}\frac{\mathrm{d}^{\beta}\gamma(t)}{\mathrm{d}t^{\beta}}$ | - | $\tan\delta = \tan\left(\frac{\pi\beta}{2}\right)$ |
| Fractional Kelvin-Voigt Liquid Model 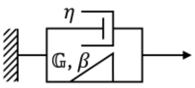  | $\sigma(t) = \eta\frac{\mathrm{d}\gamma(t)}{\mathrm{d}t} + \mathbb{G}\frac{\mathrm{d}^{\beta}\gamma(t)}{\mathrm{d}t^{\beta}}$ | $\tau = \left(\frac{\eta}{\mathbb{G}}\right)^{\frac{1}{1-\beta}}$ | $\tan\delta = \frac{\eta\omega + \mathbb{G}\omega^{\beta}\sin(\pi\beta/2)}{\mathbb{G}\omega^{\beta}\cos(\pi\beta/2)}$ |

Table S2 — Fitted quasi-properties ($\mathbb{V}$, $\mathbb{G}$) and power law exponents ($\alpha$, $\beta$) of the fractional Kelvin-Voigt Liquid model for BSM samples at 6 and 8 wt%.

| | $\alpha$ | $\mathbb{V}$ (Pa s$^{\alpha}$) | $\beta$ | $\mathbb{G}$ (Pa s$^{\beta}$) |
|---|---|---|---|---|
| 6 wt% | 1 | 0.0302 | 0.70 | 0.232 |
| 8 wt% | 1 | 0.0543 | 0.47 | 0.396 |

For the FKV-liquid model, the loss modulus is

$$G'' = \mathbb{V}\omega + \mathbb{G}\omega^{\beta}\sin(\pi\beta/2).$$

Reduction to a Scott-Blair element requires the viscous dashpot term to remain negligible throughout the experimental window, i.e.

$$\mathbb{V}\omega \ll \mathbb{G}\omega^{\beta} \sin(\pi\beta/2).$$

Enforcing this condition at the highest measured frequency, $\omega_{max}$ , gives

$$\omega_{max} \ll \left(\frac{\mathbb{G}\sin(\pi\beta/2)}{\mathbb{V}}\right)^{\frac{1}{1-\beta}}.$$

Using the fitted parameters shown in Table S2, this characteristic frequency $\omega_{max}$ is approximately $6.2 \times 10^2$ rad $s^{-1}$ for the 6 wt% sample, and $2.1 \times 10^1$ rad $s^{-1}$ for the 8 wt% sample. Consequently, for the 6 wt% sample, the condition $\omega_{max} \ll \omega_c$ is satisfied across the entire experimental window, such that $\mathbb{V}\omega \ll \mathbb{G}\omega^{\beta}$ and the viscous dashpot contribution becomes negligible. In this regime, the FKV-Liquid constitutive equation reduces to

$$\sigma(t) \approx \mathbb{G}\frac{\mathrm{d}^{\beta}\gamma(t)}{\mathrm{d}t^{\beta}},$$

which corresponds to a single spring-pot element with exponent $\beta$ and quasi-property $\mathbb{G}$, consistent with the definition in Table S1:

$$\sigma(t) = \mathbb{G}\frac{\mathrm{d}^{\beta}\gamma(t)}{\mathrm{d}t^{\beta}}.$$

The measured response is therefore governed by a single power-law, yielding an approximately frequency-independent $\tan\delta$, consistent with critical-gel like behavior within the experimental window. By contrast, for the 8 wt% sample, $\omega_c$ lies within the measured frequency range, such that the constraint $\mathbb{V}\omega \ll \mathbb{G}\omega^{\beta}$ is not satisfied at higher frequencies. Both the dashpot and fractional contributions are therefore important within the experimental window, and the full FKV-liquid description is required.

Having shown that the 6 wt% sample lies in the asymptotic regime of the FKV-Liquid model where the viscous contribution is negligible, the data were fitted with a Scott-Blair constitutive form, from which the fractional exponent $\beta$ was obtained and the corresponding frequency-independent value of the phase angle, $\tan\delta = \tan(\pi\beta/2)$, was calculated.

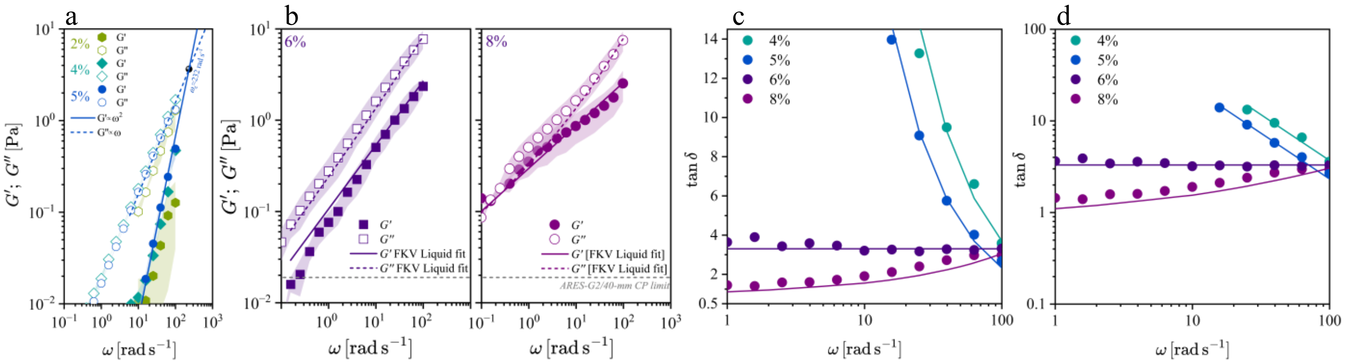

Figure S1 – (a) Linear viscoelastic response of BSM at 2, 4, and 5 wt%. Extrapolation of $G'' \sim \omega$ and $G' \sim \omega^2$ for BSM at 5 wt% yields a $G' = G''$ crossover at 232 rad s$^{-1}$ ($\tau \simeq 0.0043$s). (b) Fractional Kelvin-Voigt liquid (FKV-Liquid) model fit for 6 and 8 wt% BSM (fitted parameters are reported in Table S2). (c,d) Frequency dependence of $tan\,\delta$. Solid lines represent a single-mode Maxwell fit where $tan\,\delta = 1/(\tau\omega)$ for 4 and 5 wt% BSM, a critical gel fit for BSM at 6 wt%, where $tan\,\delta = tan(\pi\beta/2) = G''/G'$, and a FKV liquid model fit for 8 wt%, where $tan\,\delta = [\mathbb{V}\omega + \mathbb{G}\omega^{\beta}\ sin(\pi\beta/2)]/[\mathbb{G}\omega^{\beta}\ cos(\pi\beta/2)]$. Panel (c) is shown in semilogarithmic representation (logarithmic x-axis), while panel (d) is plotted on log-log axes.

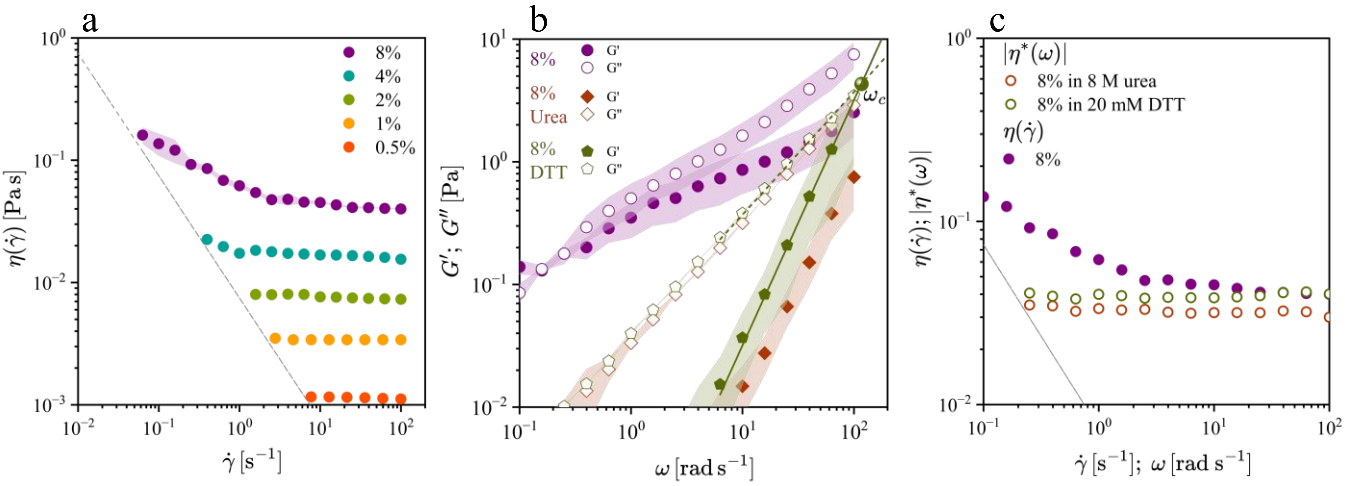

Figure S2 – (a) Steady-shear viscosity as a function of shear rate, measured using a 40 mm cone-and-plate geometry (2° CP-40 mm). The minimum rheometer sensitivity is shown as a grey dashed line. BSM samples at 8, 4, 2, 1, and 0.5 wt% were prepared in 10 mM HEPES buffer at pH 7.0. (b) Viscoelastic behavior of 8 wt% BSM in the absence or in the presence of 8 M urea, or 20 mM dithiothreitol. Shaded regions represent the standard deviations of three independent measurements. In the presence of DTT, which reduces disulfide bonds between mucin subunits, the covalent network at 8 wt% is disrupted, restoring a terminal liquid-like response with $G' \sim \omega^2$ and $G'' \sim \omega$, consistent with the Maxwell description. The corresponding relaxation time is $\tau = 8.60$ ms. In contrast, 8 wt% BSM in the presence of 8 M urea, which perturbs hydrogen bonding and weak non-covalent associations without reducing disulfide linkages, does not recover terminal behaviour of a viscoelastic liquid: while $G'' \sim \omega$ is preserved, the elastic modulus $G'$ exhibits an apparent $G' \sim \omega^{1.5}$ scaling over the accessible frequency window. (c) Steady-shear viscosity of the untreated 8 wt% BSM sample shown as a function of shear rate, compared with the complex viscosity extracted from $G'$ and $G''$, shown as a function of angular frequency, for 8 wt% BSM in the presence of either 8 M urea or 20 mM DTT.

For (b) and (c), the context of urea- and DTT-induced denaturation is different: while a small amount of DTT (20 mM) is sufficient to reduce covalent disulfide bonds, 8 M urea alone corresponds to approximately 48% w/v of the final sample. Therefore, urea-induced perturbations represent a major change in the solvent environment. The slightly lower network elasticity in 8 M urea compared to that in 20 mM DTT suggests that the residual viscous response is not controlled simply by the viscosity of the solvent but instead reflects the large perturbation of the solvent environment. This demonstrates how urea can suppress a broader set of non-covalent interactions, leading to a lower residual complex viscosity for the sample in the presence of urea.

For all steady shear or small amplitude oscillatory measurements, interfacial effects and evaporation were avoided by using a 2 vol% Tween 80 layer and a mineral oil overlay, respectively, deposited at the air-liquid interface prior to measurement.

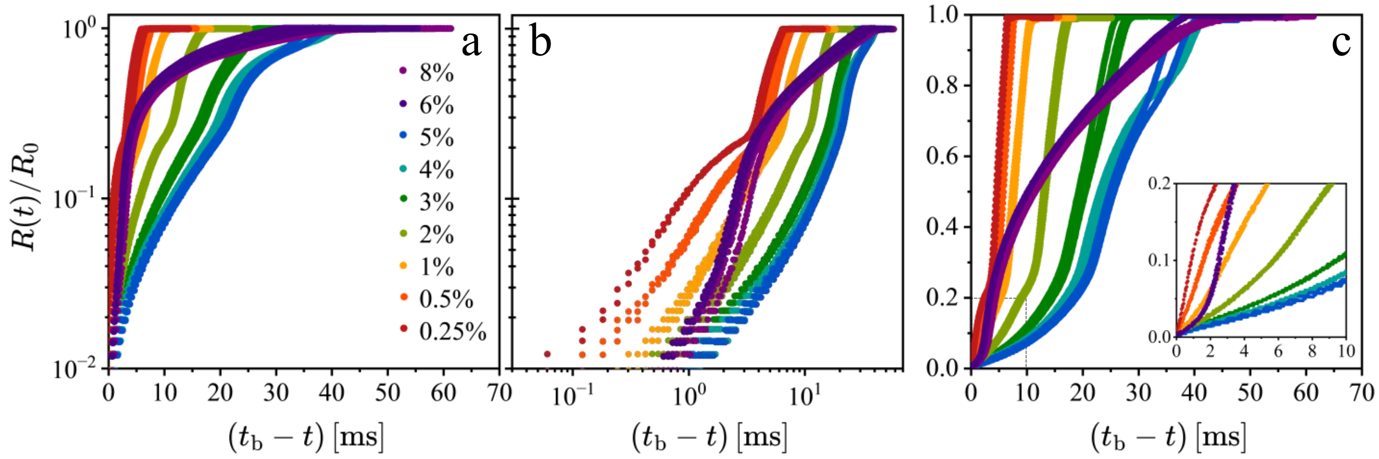

Figure S3 - Plots of the normalized filament radius $R(t)/R_0$ as a function of $(t_b - t)$, where $t_b$ is the time to breakup. For each concentration, at least three independent videos are included. In the terminal stage of the filament lifetime, replicates at a given concentration superimpose, and the trajectories for all concentrations converge as the filament approaches breakup. For completeness, data are presented using (a) log-linear, (b) log-log, and (c) linear-linear axes. The inset in (c) shows the terminal region on linear-linear axes.

**Terminal capillarity-driven thinning behavior of dilute BSM samples**

For the dilute BSM samples (Figure S3d), the capillary thinning dynamics at later stages were additionally examined in the $(t_b - t)$ representation to assess whether the filament approaches an asymptotic viscous response prior to breakup. In this limit, the minimum filament radius may be written in the general form[2]

$$\frac{R(t)}{R_0} = \frac{2X-1}{2}\left(\frac{t_b-t}{T_\infty}\right),$$

where $R_0$ is the initial filament radius, $t_b$ is the breakup time, $X$ is a geometric factor associated with the local filament shape, and $T_\infty$ is a characteristic terminal thinning timescale defined as $T_\infty = \eta_{E,\infty} R_0/\Gamma$. Here, $\eta_{E,\infty}$ denotes the asymptotic transient extensional viscosity and $\Gamma$ the surface tension. For an approximately cylindrical filament, $X \approx 1$, such that the above expression reduces to

$$\frac{R(t)}{R_0} \approx \frac{1}{2}\left(\frac{t_b-t}{T_\infty}\right).$$

Thus, if the terminal thinning regime is governed by a time-independent asymptotic extensional viscosity, $R(t)/R_0$ should vary linearly with $(t_b - t)$. On a log-log plot, this corresponds to a slope of unity, $R(t)/R_0 \sim (t_b - t)^1$. The late-time data for 0.25 and 0.5 wt% BSM samples are consistent with this scaling, supporting the interpretation of a terminal response in the final stage of thinning (see Figure S3d). Assuming $X \approx 1$, the terminal thinning regime was fitted in the interval $10^{-1} \leq (t_b - t) \leq 10^0$ [ms] using the form $R(t)/R_0 = A(t_b - t)$, where $A$ is the fitted prefactor and the slope was fixed to unity.

Comparison with the asymptotic expression above gives $A = 1/2T_\infty$, so that $T_\infty = 1/2A$ and hence $\eta_{E,\infty} = \Gamma T_\infty/R_0 = \Gamma/2AR_0$. Using $R_0 = 0.635$ mm, and $\Gamma = 61.7$ mN m$^{-1}$, the fitted prefactor for the 0.5 wt% BSM sample $A = 63.5655$ s$^{-1}$ gives $T_\infty = 7.87 \times 10^{-3}$ s and $\eta_{E,\infty} = 0.77$ Pa s. Likewise, for the 0.25% BSM sample, $A = 115.525$ s$^{-1}$, giving $T_\infty = 4.33 \times 10^{-3}$ s and $\eta_{E,\infty} = 0.42$ Pa s. These asymptotic extensional viscosities obtained close to filament breakup of the dilute BSM samples are consistent with the corresponding apparent transient extensional viscosity values for the low concentration fluids shown in Figure S6, which may indicate that the dilute samples approach a terminal finite-extensibility dominated thinning regime prior to breakup.

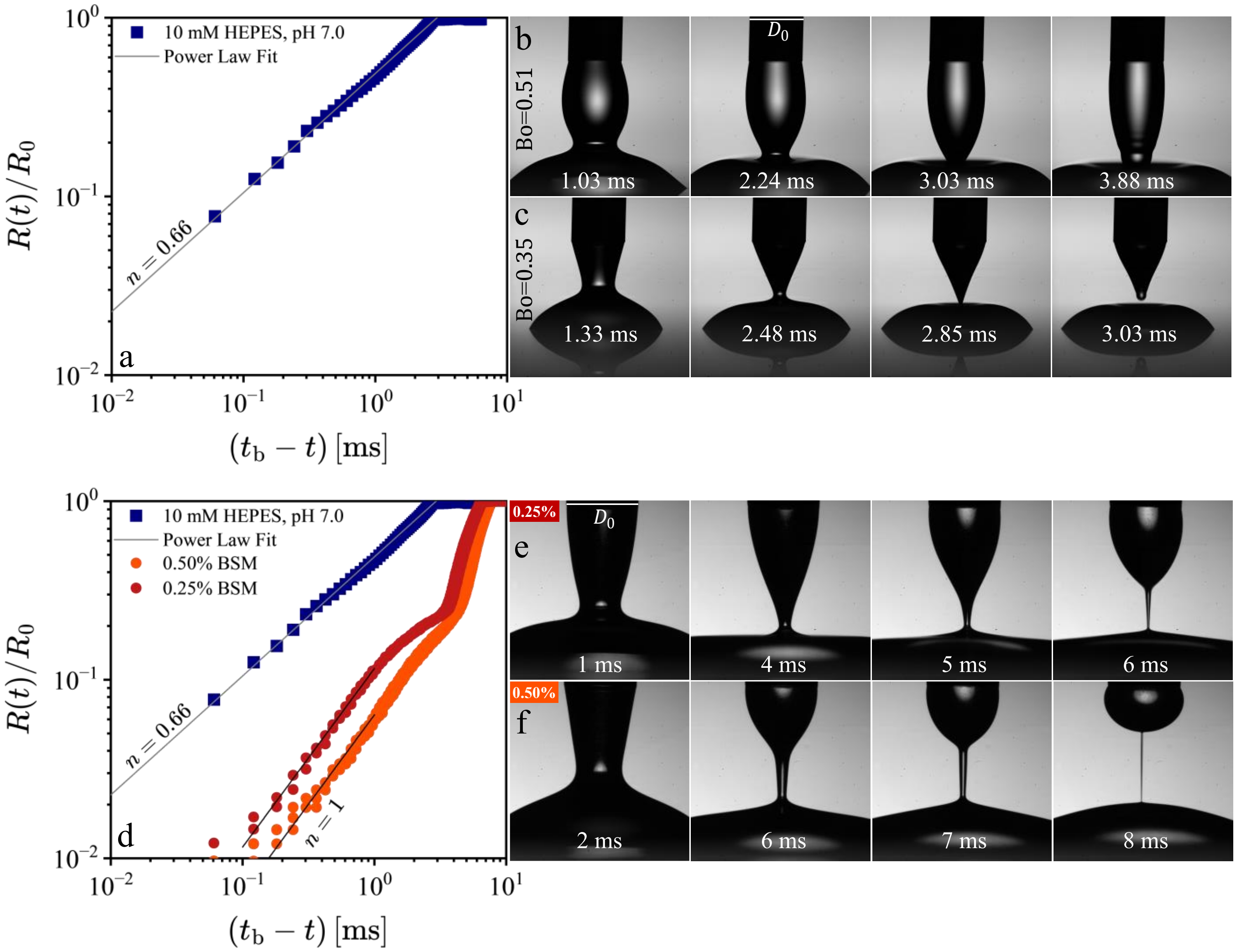

Figure S4 – Dripping-on-substrate measurements with $D_0 = 2R_0 = 1.27$ mm (a) for the solvent used in this work [10 mM HEPES buffer at pH 7.0 (blue)]. The data for 10 mM HEPES buffer at pH 7.0 are shown for the profile extracted from images performed at a Bond number $Bo = 0.35$. Snapshots of 10 mM HEPES buffer at pH 7.0 at a separation distance that yield Bond numbers of (b) $Bo = 0.51$ and (c) $Bo = 0.35$. The conical shape is most clearly observed at smaller needle-substrate separation distances $H$, corresponding to lower Bond numbers ($Bo = R_0 H\rho g/\Gamma$), as gravitational effects are reduced relative to capillarity at the moment the pendant drop touches the substrate. For the buffer, the Ohnesorge number $Oh = \eta_0/\sqrt{\rho\Gamma R_0}$ is $Oh \approx 4.4 \times 10^{-3}$, placing the system well within the inertio-capillary regime. The time to breakup ($t_b$) for HEPES buffer is 2.78 ms. (d) Normalized minimum filament radius $R(t)/R_0$ as a function of $t_b - t$ for 0.25 and 0.5 wt% BSM. In the final stage of thinning, both samples approach $R(t)/R_0 \sim (t_b - t\,)^1$ as indicated by the slope $n = 1$, consistent with a viscosity-dominated terminal thinning regime. (e-f) Corresponding high-speed image sequences for (e) 0.25 wt% ($t_b = 6.24$ ms) and (f) 0.5% BSM ($t_b = 8.11$ ms).

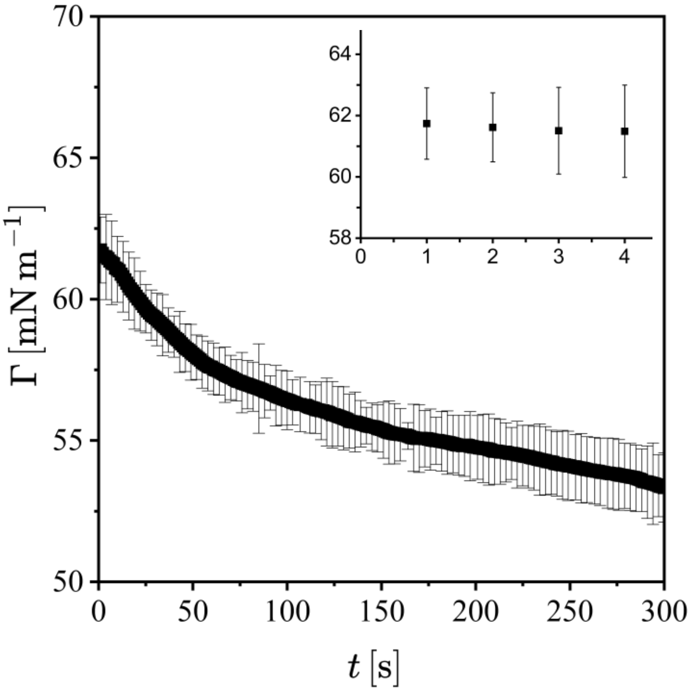

Figure S5 – Surface tension (Γ, mN m$^{-1}$) of BSM at 2 wt% as a function of time (s). Mean ± standard deviation from triplicate measurements is shown. Across all concentrations (0.25-8 wt%), the initial surface tension is indistinguishable within experimental resolution. Deviations only emerge at longer timescales, beyond the temporal window relevant to DoS measurements (< 1 s). Inset: variation of surface tension over 1-4 s, highlighting negligible early-time variations.

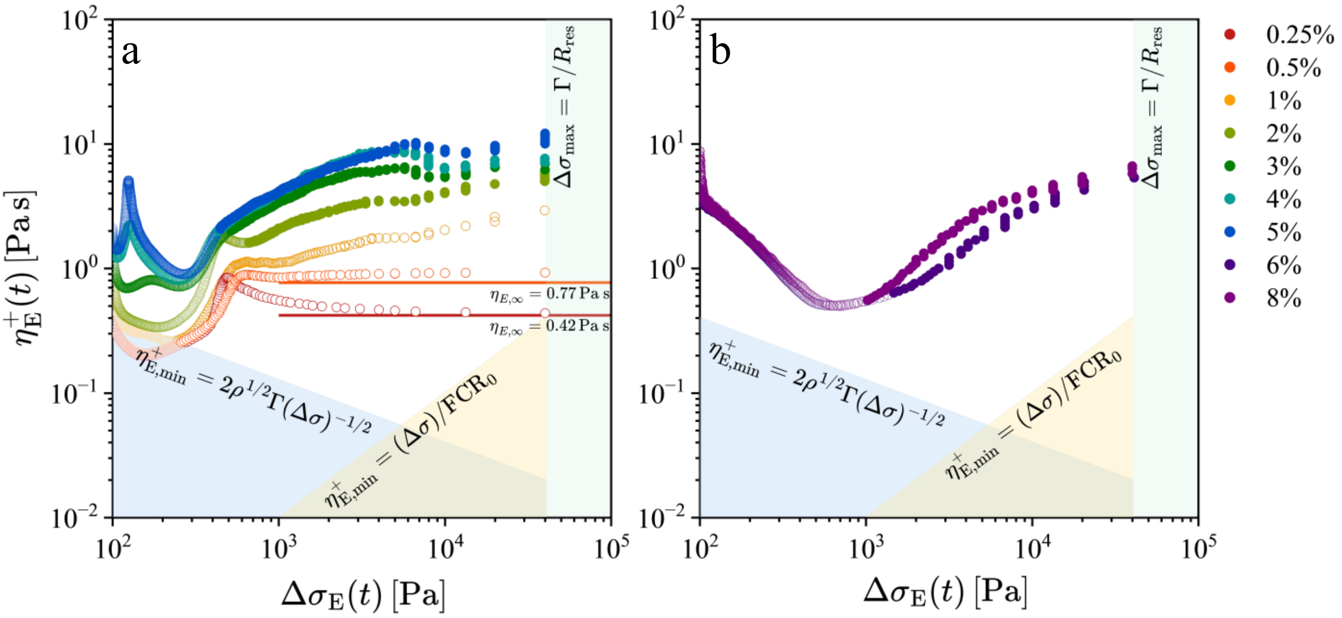

Figure S6 - Apparent transient extensional viscosity plotted as a function of the time-evolving extensional stress difference for BSM dispersions spanning 1 to 8 wt%. Limit lines, as discussed in reference 13 of the manuscript, are drawn using $\rho = 1080$ kg m$^{-3}$, $\Gamma = 61.7$ mN m$^{-1}$, $R_0 = 0.635$ mm, $R_{res} = 1$ pixel $= 1.51 \times 10^{-6}$ m, $FCR_0 = f_s \, ln \, m_0$, with $f_s = 16500$ s$^{-1}$ and $m_0 = R_0/R_{res} = 430.46$. The horizontal lines shown in (a) indicate the asymptotic transient extensional viscosities $\eta_{E,\infty}$, as previously discussed (see the discussion on page S-8).

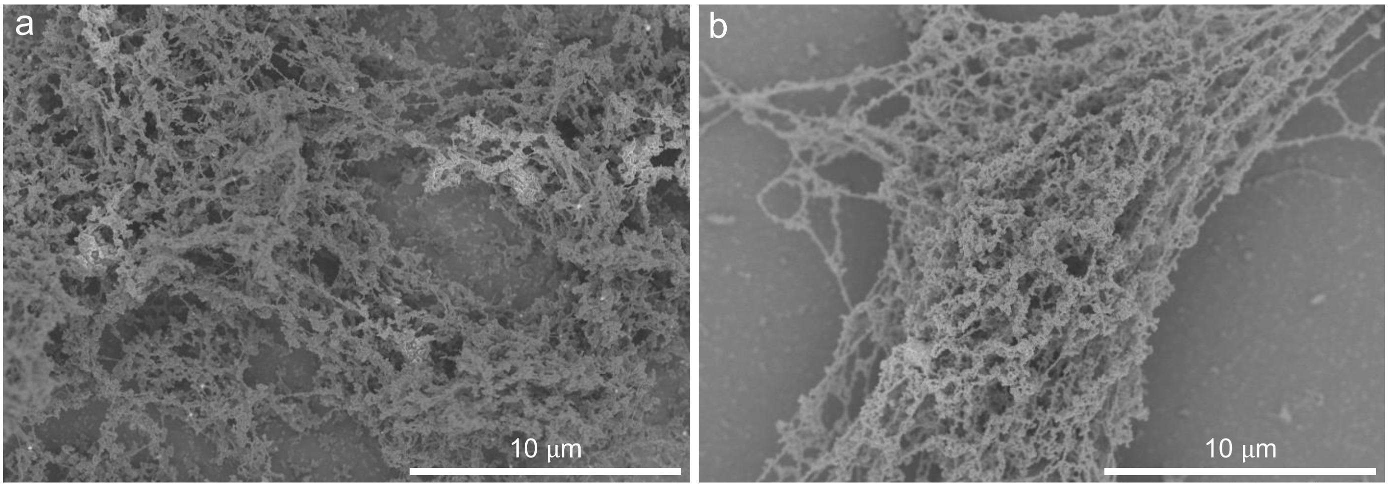

Figure S7 - Representative Scanning Electron Microscopy images of BSM samples prepared at (a) 6 wt% and (b) 5 wt%. Scales bars represent 10 μm.

Table S3 — Measured surface tension using a pendant drop tensiometer (OCA 25, DataPhysics Instruments, Germany). Drops (22 μL) were formed at the tip of a 1.65 mm DS500 GT syringe, and the liquid-air surface tension was obtained by fitting the drop profile to the Young-Laplace equation using dpiMAX software. A density of 1080 kg m$^{-3}$ was assumed in the fitting procedure. Measurements were performed in triplicate for each concentration; the reported value include the standard deviation. The initial surface tension was used in all capillarity-driven thinning analyses, as filament thinning occurs on timescales much shorter than those required for interfacial equilibration.

| BSM concentration (wt%) | Surface tension (mN m$^{-1}$) |
|---|---|
| 0.25-8 wt% | $61.74 \pm 2.3$ |

Table S4 — Reference Ohnesorge number ($Oh = \eta_0/\sqrt{\rho \Gamma R_0}$) for the BSM concentrations investigated. Here, $\rho = 1080$ kg m$^{-3}$, $\Gamma = 61.74$ mN m$^{-1}$, $R_0 = 0.635$ mm. The reported value of $\eta_0$ is an estimate from the low-shear viscosity as the zero-shear plateau was not directly resolved in steady shear.

| BSM concentration (wt%) | $\eta_0$ / Pa s | $Oh$ |
|---|---|---|
| 0.25 | 0.0010 | 0.005 |
| 0.5 | 0.0012 | 0.006 |
| 1 | 0.0035 | 0.017 |
| 2 | 0.0079 | 0.038 |
| 4 | 0.0225 | 0.109 |
| 8 | 0.1604 | 0.780 |

Table S5 — Comparison between relaxation times ($\tau$) obtained from SAOS estimated from the extrapolated crossover frequency for 2-5 wt% (and from $\tau = (\mathbb{V}/\mathbb{G})^{1/(1-\beta)}$ at 8 wt%), and the effective relaxation times extracted from elastocapillary (EC) thinning regimes in DoS extensional experiments ($\tau_E$) for BSM across concentrations. The time to breakup ($t_b$) for each fluid is also shown.

| BSM concentration (wt%) | $\tau$ (SAOS) / ms | $\tau_E$ (DoS) / ms | $t_b$ / ms |
|---|---|---|---|
| 0.25 | - | - | $6.24 \pm 0.3$ |
| 0.5 | - | - | $8.11 \pm 0.2$ |
| 1 | - | $0.53 \pm 0.08$ | $12.2 \pm 0.6$ |
| 2 | 1.6 | $1.34 \pm 0.05$ | $18.2 \pm 0.5$ |
| 3 | - | $2.4 \pm 0.08$ | $28.4 \pm 1.4$ |
| 4 | 2.7 | $3.0 \pm 0.26$ | $39.7 \pm 1.9$ |
| 5 | 4.3 | $2.8 \pm 0.15$ | $41.5 \pm 0.3$ |
| 6 | critical gel | $0.34 \pm 0.02$ | $42.8 \pm 2.5$ |
| 8 | 23.6 | $0.33 \pm 0.05$ | $47.1 \pm 3.1$ |

References

(1) Song, J.; Holten-Andersen, N.; McKinley, G. H. Non-Maxwellian viscoelastic stress relaxations in soft matter. *Soft Matter* **2023**, *19* (41), 7885–7906. DOI: 10.1039/d3sm00736g.
(2) Dinic, J.; Sharma, V. Macromolecular relaxation, strain, and extensibility determine elastocapillary thinning and extensional viscosity of polymer solutions. *PNAS* **2019**, *116* (8), 8766–8774. DOI: 10.1073/pnas.1820277116.
